# Oral and injected tamoxifen alter adult hippocampal neurogenesis in female and male mice

**DOI:** 10.1101/2021.09.30.462662

**Authors:** Bryon M. Smith, Angela I. Saulsbery, Patricia Sarchet, Nidhi Devasthali, Dalia Einstein, Elizabeth D. Kirby

## Abstract

Inducible Cre recombinase facilitates temporal control of genetic recombination in numerous transgenic model systems, a feature which has made it a popular tool for studies of adult neurogenesis. One of the most common forms of inducible Cre, CreER^T2^, requires activation by the synthetic estrogen tamoxifen (TAM) to initiate recombination of LoxP-flanked sequences. To date, most studies deliver TAM via intraperitoneal injection. But the introduction of TAM-infused commercial chows has recently expanded the possible modes of TAM delivery. Despite the widespread use of TAM-inducible genetic models in adult neurogenesis research, the comparative efficiency and off-target effects of TAM administration protocols is surprisingly infrequently studied. Here we compare a standard, 5 day TAM injection regimen with voluntary consumption of TAM-infused chow. First, we used adult NestinCreER^T2^;Rosa-LoxP-STOP-LoxP-EYFP reporter mice to show that 2 weeks of TAM chow and 5 days of injections led to LoxP recombination in a similar phenotypic population of neural stem and progenitor cells in the adult dentate gyrus. However, TAM chow resulted in substantially less overall recombination than injections. TAM administration also altered adult neurogenesis, but in different ways depending on administration route: TAM injection disrupted neural progenitor cell proliferation 3 weeks after TAM, whereas TAM chow increased neuronal differentiation of cells generated during the diet period. These findings provide guidance for selection of TAM administration route and appropriate controls in adult neurogenesis studies using TAM-inducible Cre mice. They also highlight the need for better understanding of off-target effects of TAM in other neurological processes and organ systems.

## Introduction

Adult neurogenesis is a widely conserved process among mammalian species in which resident neural stem cells generate new neurons that integrate into mature circuitry throughout the lifespan. Evidence that neurogenesis in the adult rodent dentate gyrus of the hippocampus supports memory and affect regulation has spurred interest in both its natural functions across species, as well as its potential therapeutic applications in humans (for reviews, see (McAvoy and Sahay, 2017; Miller and Sahay, 2019; Toda et al., 2019).

The study of time- and tissue-specific phenomena, such as adult hippocampal neurogenesis, has been accelerated by the introduction of several inducible methods of gene expression manipulation. Tamoxifen (TAM)-inducible Cre mouse lines (e.g. CreER^T2^) have proven to be particularly valuable tools for inducible manipulation of gene expression in the adult neurogenic lineage. In these models, Cre recombinase is fused to a mutated estrogen receptor (ER^T2^) which retains Cre in the cytoplasm until the receptor binds its ligand, the synthetic estrogen tamoxifen (TAM) (Feil et al., 1997; Danielian et al., 1998). TAM binding allows translocation of the mutated estrogen receptor and its fused Cre enzyme to the nucleus, where Cre can induce recombination of LoxP-flanked genetic sequences. Use of a tissue-specific promoter can further constrain CreER^T2^ expression to specific cell classes. For example, neural stem and progenitor cells are commonly targeted using GLAST, Nestin, SOX2, or GFAP promoters (to name a few common versions) (Semerci and Maletic-Savatic, 2016).

In adult neurogenesis studies using CreER^T2^ mediated LoxP recombination, two dominant approaches to selecting controls have emerged. One approach compares mice of different genotypes (Cre+/- or LoxP+/-), all of which are treated with TAM. A second approach is to compare mice of the same genotype treated with TAM versus those treated with vehicle. This latter comparison is predicated on TAM itself being inert, aside from its ability to induce CreER^T2^ translocation. Yet, gonadal steroid hormones, including estrogens, can modulate brain processes such as neurogenesis and cell survival (Jorgensen and Wang, 2020).

The effect of TAM on adult neurogenesis is generally understudied and the few existing studies have yielded mixed results. For example, (Rotheneichner et al., 2017) found no inherent effects of TAM on cell proliferation and fate in the dentate gyrus of the adult mouse hippocampus, but a more recent study suggested long-lasting suppression of hippocampal cell proliferation in juvenile mice (Lee et al., 2020). Given the widespread use of TAM-inducible models in adult neurogenesis, it is imperative to understand how TAM itself (independent of transgene expression) affects adult neurogenic processes.

Beyond the selection of controls, TAM administration route is another important potential design variable in studies using CreER^T2^ mouse models. Intraperitoneal injections are by far the most common method of exposing mice to TAM in adult neurogenesis studies to date. However, administration of TAM through voluntary consumption of custom laboratory chow is a promising alternative that might reduce injection-related stress and researcher hands-on time. TAM-infused chows have been used successfully in cardiac research (Kiermayer et al., 2007), and may be applicable to other adult tissues as well (Yoshinobu et al., 2021). It is unknown how TAM chow compares to TAM injection as regards recombination efficiency and optimal dosing schedule for adult neurogenesis research.

Here, we compare TAM administration by injection and chow, both in terms of the specificity and efficiency of LoxP recombination and the inherent effects of TAM treatment itself on adult neurogenesis. Using a CreER^T2^ model targeted to neural stem and progenitor cells (NSPCs) by a Nestin promoter, our findings suggest that TAM chow can induce LoxP recombination in hippocampal NSPCs with similar specificity as TAM injection, but that TAM chow results in much lower recombination efficiency than injections, likely due to chow avoidance and therefore lower TAM exposure. We also show that both TAM injection and TAM chow paradigms have inherent, though different, effects on adult neurogenesis, underscoring the need to compare TAM-treated experimental animals to genetic controls similarly treated with TAM, regardless of route of administration.

## Results

### TAM chow results in weight loss and less total TAM exposure compared to TAM injection

To compare TAM-induced LoxP recombination in NSPCs between injection and chow-fed methods, we crossed NestinCreER^T2^ mice (Lagace et al., 2007) with Rosa26-LoxP-STOP-LoxP-EYFP mice (Srinivas et al., 2001). At 7-9 weeks of age, NestinCreER^T2+/-^;Rosa26-LoxP-STOP-LoxP-EYFP^+/-^ male and female mice were assigned randomly to either injection or chow groups. Injected mice received a standard 5d injection regimen and were perfused on day 7 or 14 after the start of injections (Fig 1A). TAM chow-fed mice were given free access to TAM chow for 10 or 14d before perfusion (Fig 1A). Both male and female TAM chow-fed mice showed significant weight loss compared to TAM-injected mice (Fig 1B,C). TAM chow-fed mice consumed very little of the chow for the first ∼5 days (Fig 1D), a time period which coincided with the greatest weight loss in these mice. Mice appeared active and well-groomed throughout the experiment. By day 6-8, male and female mice significantly increased their chow consumption, coinciding with stabilization of their body weights. Cumulative TAM exposure was 1.23-1.57 fold higher in injected mice than chow-fed mice (Fig 1E). Together, these data suggest that mice find TAM chow aversive, resulting in less TAM exposure over 14d than from 5d of standard injections.

**Figure 1.**
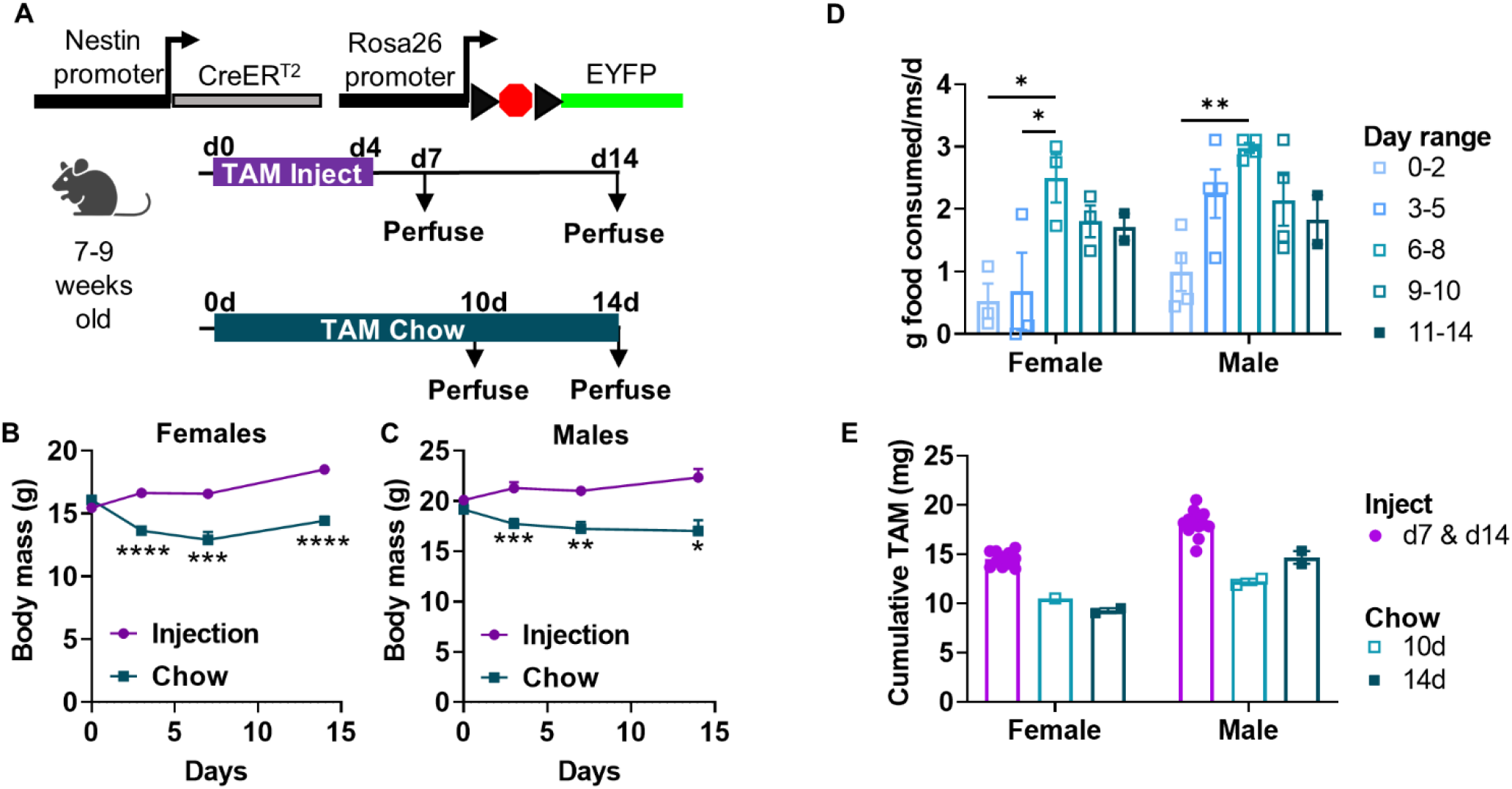
TAM chow reduces body mass and results in less TAM exposure compared with TAM injection. A) Experimental model and timeline. Adult Nestin-CreER^T2+/-^;Rosa26-LoxP-STOP-LoxP-EYFP^+/-^ mice were either injected with TAM (IP) or voluntarily fed TAM chow. B, C) Body mass (in g) of female (Day: *F*_*(1*.*637, 24*.*56)*_ = 8.314, *p* = 0.0029; TAM: *F*_*(1, 20)*_ = 45.35, *p* < 0.0001; Day x TAM interaction: *F*_*(3, 45)*_= 23.42, *p* < 0.0001) (B) and male (Day: *F*_*(0*.*5896, 9*.*630)*_ = 1.338, *p* = 0.2412; TAM: *F*_*(1, 23)*_ = 21.96, *p* = 0.0001; Day x TAM interaction: *F*_*(3, 49)*_= 10.40, *p* < 0.0001) (C) mice during and after TAM treatment. Mean ± SEM of n = 7-11 mice. D) TAM chow consumption (g of food) per mouse per day, averaged within cage (Sex: *F*_*(1, 22)*_ = 6.139, *p* = 0.0214; Day: *F*_*(4, 22)*_ = 8.310, *p* = 0.0003; Interaction: *F*_*(4, 22)*_= 1.159, *p* = 0.3558). Mean ± SEM of n = 2-4 cages shown. E) Cumulative TAM exposure (mg of TAM) in injected and chow-fed mice (Sex: *F*_*(1, 25)*_ = 39.49, *p* < 0.0001; TAM: *F*_*(2, 25)*_= 51.35, *p* < 0.0001; Interaction: *F*_*(2, 25)*_ = 2.626, *p* = 0.0922). Mean ± SEM shown from n = 11-13 mice (injection) or n = 1-2 cages (chow). \**P* < 0.05, \*\**P* < 0.01, \*\*\**P* < 0.001, \*\*\*\**P* < 0.0001, as determined by Sidak’s multiple comparisons test following a mixed-effects analysis (B-C) or 2-way ANOVA (D-E).

### TAM chow results in lower recombination efficiency then TAM injections

We next compared total EYFP+ cell number in the subgranular zone of the DG to determine the relative efficacy of TAM-induced Rosa26-LoxP-STOP-LoxP-EYFP recombination in these mice. As expected based on previous studies with this line of NestinCreER^T2^ mice (Lagace et al., 2007; Sun et al., 2014; Dause and Kirby, 2020), EYFP expression filled cell bodies and was strongly localized to the SGZ in both TAM injected and TAM chow-fed mice (Fig 2A). 10d and 14d TAM chow-fed mice had 2.54-fold and 3.31-fold fewer EYFP+ cells compared to injected mice perfused 14d after the start of injections (Fig 2A,B). We next quantified the efficiency of EYFP expression specifically in GFAP+/SOX2+ radial glia-like NSCs (RGL-NSCs) and GFAP-/SOX2+ intermediate progenitor cells (IPCs) (Fig 2C). TAM injected mice showed EYFP expression in slightly less than half of RGL-NSCs (47.55 ± 6.34% and 47.18 ± 4.63% at 7d and 14d after injection start) while TAM chow fed mice showed EYFP expression in less than a quarter of RGL-NSCs (23.61 ± 6.72% and 13.80 ± 2.53% at 10d and 14d of chow) (Fig 2D). Among IPCs, TAM injected mice showed EYFP expression in 39.86 ± 10.00% and 55.91 ± 7.65% of IPCs at 7d and 14d after injection start. TAM chow fed mice again showed significantly less EYFP expression among IPCs (29.89 ± 12.72% and 24.55 ± 5.15% at 10d and 14d of chow) (Fig 2D). Separate analysis of males and females yielded similar results in each sex (Fig 3A-B). Together, these findings suggest lower, but detectable, LoxP-recombination following 10-14d of TAM chow feeding compared to a standard 5d injection regimen.

**Figure 2.**
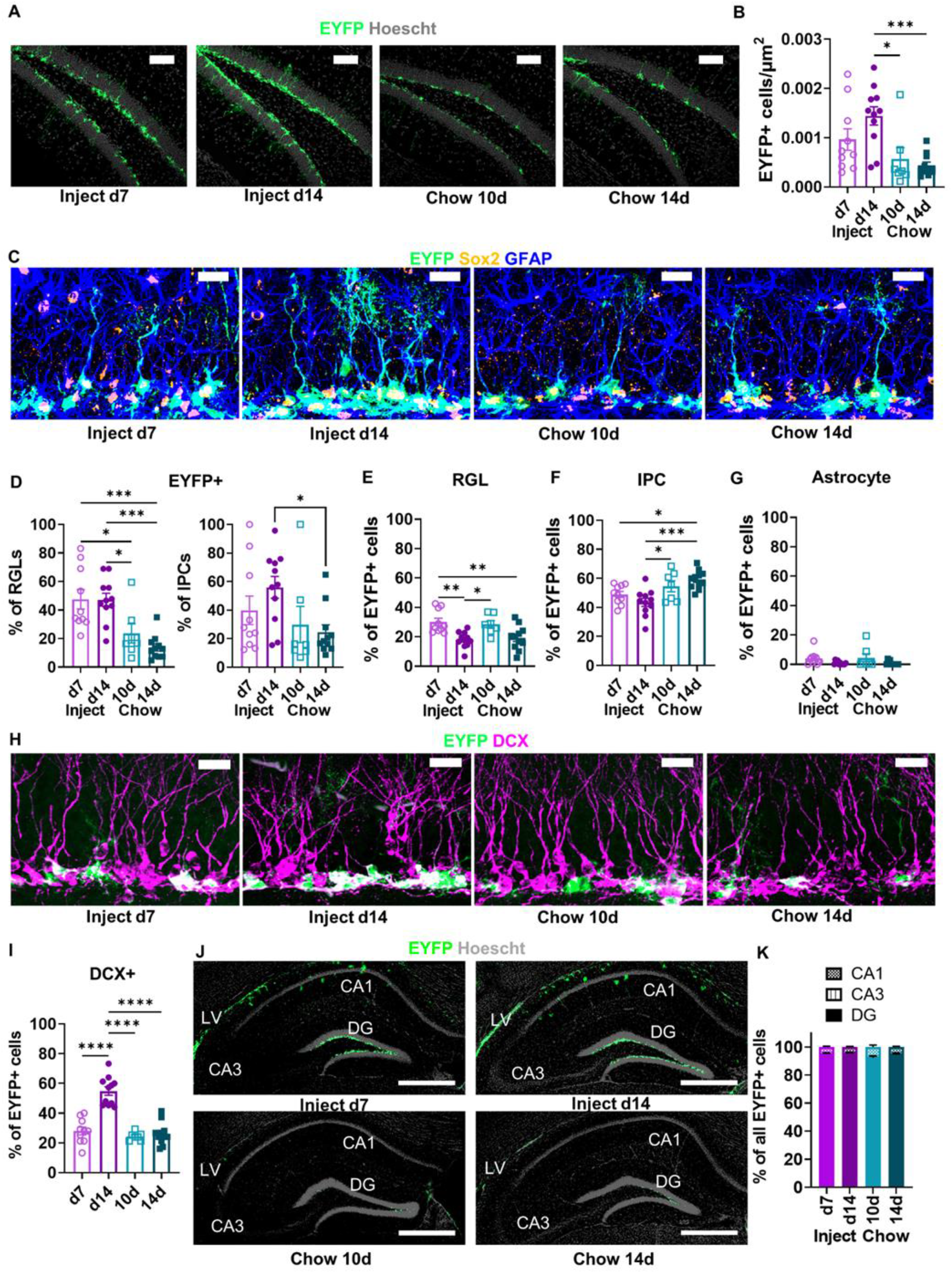
Recombination efficiency is higher following TAM injection than TAM diet. A) Representative images of EYFP+ labelling in the DG. Scale = 100 μm. B) Density of EYFP+ cells (cells/μm^2^) in the DG (ordinary one-way ANOVA, *F*_*(3, 35)*_ = 6.885, *p* = 0.0009). Mean ± SEM of n = 7-11 mice. C) Representative images of EYFP, SOX2, and GFAP labelling in the DG. Scale = 20 μm. D) The percent of phenotypic GFAP+/SOX2+ RGL-NSCs and GFAP-/SOX2+ IPCs that were EYFP+ (RGLs: ordinary one-way ANOVA, *F*_*(3, 35)*_ = 11.43, *p* < 0.0001; IPCs: ordinary one-way ANOVA, *F*_*(3, 35)*_ = 2.788, *p* = 0.0550). Mean ± SEM of n = 7-11 mice. E) The percent of EYFP+ cells that showed RGL-NSC phenotype (ordinary one-way ANOVA, *F*_*(3, 35)*_ = 7.531, *p* = 0.0005). Mean ± SEM of n = 7-11 mice. F) The percent of EYFP+ cells that showed IPC phenotype (ordinary one-way ANOVA, *F*_*(3, 35)*_ = 7.799, *p* = 0.0004). Mean ± SEM of n = 7-11 mice. G) The percent of EYFP+ cells that showed GFAP+ stellar astrocyte phenotype (ordinary one-way ANOVA, *F*_*(3, 35)*_ = 2.465, *p* = 0.0784). Mean ± SEM of n = 7-11 mice. H) Representative images of EYFP and DCX labelling in the DG. Scale = 20 μm. I) The percent of EYFP+ cells that were DCX+ (ordinary one-way ANOVA, *F*_*(3, 35)*_ = 34.97, *p* < 0.0001). Mean ± SEM of n = 7-11 mice. J) Representative images of EYFP labelling throughout the hippocampus. Scale = 500 μm. K) Percent of hippocampal EYFP+ cells found in the CA1, CA3, and DG (2-way repeated measures ANOVA; TAM x area: *F*_*(6, 70)*_ = 3.338, *p* = 0 0060; TAM: *F*_*(3, 35)*_ = 1.051, *p* = 0.3822; area: *F*_*(1*.*186, 41*.*50)*_= 20157, *p* < 0.0001; subject: *F*_*(35, 70)*_ = 6.492e-007, *p >* 0.9999). Mean ± SEM of n = 7-11 mice. \**P*< 0.05, \*\**P*< 0.01, \*\*\**P*< 0.001, \*\*\*\**P*< 0.0001, as determined by Tukey’s multiple comparisons test.

**Figure 3.**
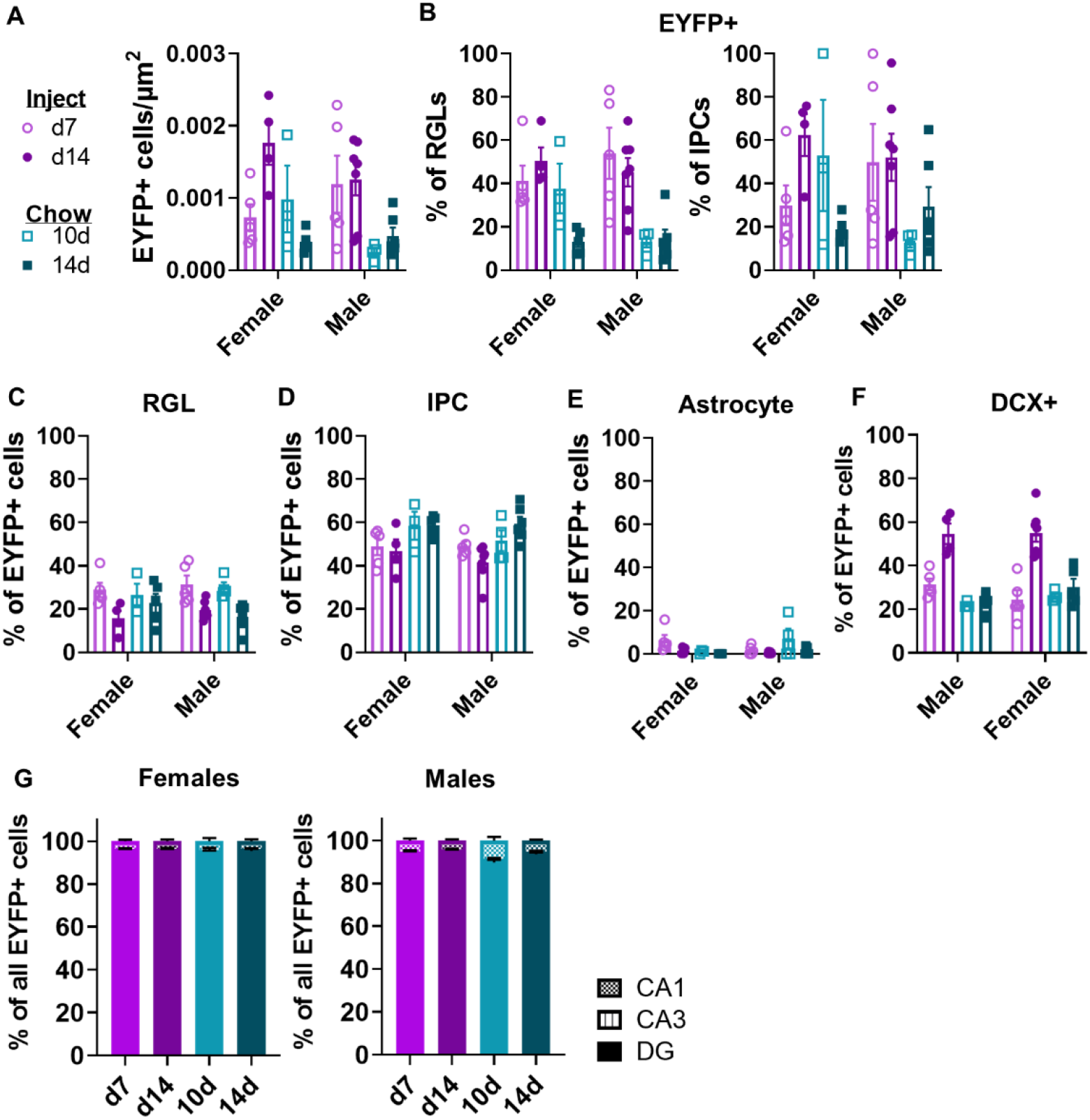
Recombination efficiency and labeled cell types do not depend on sex. A) Density of EYFP+ cells (cell/μm^2^) in the DG by sex (2-way ANOVA; Interaction: *F*_*(3, 31)*_ = 2.335, *p* = 0.0931; sex: *F*_*(1, 31)*_ = 0.9785, *p* = 0.3302; TAM: *F*_*(3, 31)*_ = 8.017, *p* = 0.0004). Mean ± SEM of n = 3-7 mice. B) The percent of phenotypic GFAP+/SOX2+ RGL-NSCs (2-way ANOVA; Interaction: *F*_*(3, 31)*_ = 2.003, *p* = 0.1341; sex: *F*_*(1, 31)*_ = 0.5780, *p* = 0.4528; TAM: *F*_*(3, 31)*_ = 11.87, *p* < 0.0001) and GFAP-/SOX2+ IPCs (2-way ANOVA; Interaction: *F*_*(3, 31)*_ = 2.152, *p* = 0.1138; sex: *F*_*(1, 31)*_ = 0.3609, *p* = 0.5524; TAM: *F*_*(3, 31)*_ = 3.040, *p* = 0.0436) that were EYFP+, with sex on the horizontal axis. Mean ± SEM of n = 3-7 mice. C) The percent of EYFP+ cells that showed RGL-NSC phenotype, analyzed separately by sex (2-way ANOVA; Interaction: *F*_*(3, 31)*_ = 1.147, *p* = 0.3457; sex: *F*_*(1, 31)*_ = 0.1478, *p* = 0.7032; TAM: *F*_*(3, 31)*_ = 7.335, *p* = 0.0007). Mean ± SEM of n = 3-7 mice. D) The percent of EYFP+ cells that showed IPC phenotype, analyzed separately by sex (2-way ANOVA; Interaction: *F*_*(3, 31)*_ = 0.4952, *p* = 0.6883; sex: *F*_*(1, 31)*_ = 1.078, *p* = 0.3072; TAM: *F*_*(3, 31)*_ = 6.650, *p* = 0.0013). Mean ± SEM of n = 3-7 mice. E) The percent of EYFP+ cells that showed GFAP+ stellar astrocyte phenotype, analyzed separately by sex (2-way ANOVA; Interaction: *F*_*(3, 31)*_ = 2.982, *p* = 0.0464; sex: *F*_*(1, 31)*_= 0.1198, *p* = 0.7316; TAM: *F*_*(3, 31)*_ = 2.411, *p* = 0.0857). Mean ± SEM of n = 3-7 mice. F) The percent of EYFP+ cells that showed DCX+ immature neuron phenotype, analyzed separately by sex (2-way ANOVA; Interaction: *F*_*(3, 31)*_ = 1.580, *p* = 0.2141; sex: *F*_*(1, 31)*_ = 0.2172, *p* = 0.6444; TAM: *F*_*(3, 31)*_ = 33.62, *p* < 0.0001). Mean ± SEM of n = 3-7 mice. G) Percent of hippocampal EYFP+ cells found in the CA1, CA3, and DG in male (2-way repeated measures ANOVA; group x region: *F*_*(6, 36)*_ = 6.259, *p* = 0.0001; group: *F*_*(3, 18)*_= 1.636, *p* = 0.2162; region: *F*_*(1*.*303, 23*.*46)*_ = 14034, *p* < 0.0001; subject: *F*_*(18, 36)*_ = 3.413e-007, *p >* 0.9999) and female (2-way repeated measures ANOVA; group x region: *F*_*(6, 26)*_= 0.2057, *p* = 0.9719; group: *F*_*(3, 13)*_ = 1.090, *p* = 0.3880; region: *F*_*(1*.*193, 15*.*51)*_ = 12616, p < 0.0001; subject: *F*_*(13, 26)*_ ^=^ 1.595e-006, p > 0.9999) mice at d7 and d14 dpi and 10d and 14d after TAM chow initiation. Mean ± SEM of n = 3-5 female mice and 4-7 male mice. \**P*< 0.05, \*\**P*< 0.01, \*\*\**p*< 0.001, \*\*\*\**P*<0.0001, as determined by Sidak’s multiple comparisons test (A-F) or Tukey’s multiple comparisons test (G).

### TAM chow and TAM injections show similar recombination specificity

We next quantified the cell phenotype of EYFP+ cells in the DG. At day 7 after TAM injection start, 30.11 ± 2.49% of DG EYFP+ cells were phenotypic RGL-NSCs (Fig 2E). Mice fed TAM chow for 10d showed a similar fraction of EYFP+ cells that were phenotypic RGL-NSCs (28.48 ± 2.48%). By 14d after TAM injection start, the percent of EYFP+ cells that were RGL-NSCs dropped significantly compared to 7d to 17.95 ± 1.61%. This decrease in proportion of EYFP+ cells that are RGL-NSCs is consistent with differentiation of the recombined, EYFP+ population over time. In the 14d chow group, 19.28 ± 2.47% of EYFP+ cells were phenotypic RGL-NSCs, a fraction that was slightly but significantly lower than the 7d injection group. The percent of EYFP+ cells showing IPC phenotype were similar at 7d and 14d after injection start (48.99 ± 2.73% and 43.44 ± 2.73%) (Fig 2F). However, TAM chow-fed mice showed slightly, though significantly, higher percentages of EYFP+ cells that were phenotypic IPCs than TAM-injected mice (10d: 54.54 ± 3.82%; 14d: 59.17 ± 1.91%). As expected, very few EYFP+ cells showed an astrocytic phenotype in all groups (d7 inject: 4.00 ± 1.45%; d14 inject: 0.83 ± 0.36%; 10d chow: 4.40 ± 2.76%; 14d chow: 0.61 ± 1.38%) (Fig 2G). We also used co-labeling of doublecortin (DCX) with EYFP to quantify the portion of EYFP+ cells that were phenotypic neuroblasts/immature neurons. DCX co-labeling increased significantly from 7d to 14d after TAM injection start (28.00 ± 2.61% to 54.91 ± 2.90% of EYFP+ cells co-labeled for DCX), again consistent with the maturation of the recombined, EYFP+ cell population over time. Both TAM chow-fed groups showed similar percentages of EYFP+ cells co-labeling for DCX as the 7d injection group (10d: 24.44 ± 1.05%; 14d: 26.07 ± 2.29%). Separate analysis of males and females yielded similar results in each sex (Fig 3C-F).

To assess ectopic recombination in non-neurogenic regions of the hippocampus, we quantified EYFP+ cells in CA1 and CA3. In all groups, EYFP+ cells were detectable in CA1 and CA3 but the rates of EYFP expression were low, with >92% of all EYFP+ cells being located in the DG (Fig 2J-K). Groups did not significantly differ from each other in EYFP localization to the DG and separate analysis of males and females yielded similar results in each sex (Fig 3G). All together, these data suggest that both TAM injection and TAM chow drive recombination predominantly in NSPCs with no notable differences in their specificity. They also suggest that 10d-14d of chow exposure leads to a recombined population that is most phenotypically similar to that found 7d after TAM injection start.

### TAM injection leads to a long-term suppression of cell proliferation

We next asked whether TAM exposure itself disrupts physiological adult hippocampal neurogenesis. Adult wildtype C57BL6/J mice were injected with TAM or matched volumes of vehicle (oil) for 5d (Fig 4A). Mice received daily injections of bromodeoxyuridine (BrdU) to label dividing cells coincident with the last 3 days of TAM. Mice were perfused 27d after the first TAM injection (21d after the last TAM/BrdU injections), 2h after a single injection of ethynyldeoxyuridine (EdU) to label acutely proliferating cells. Body weights of female and male mice were not altered by 5d TAM injection compared to vehicle injection (Female: 17.38 ± 0.36 g veh vs. 17.80 ± 0.38 g TAM; Male: 21.93 ± 0.72 g veh vs 22.38 ± 0.46 g TAM) (Fig 4B,C). However, after 21d recovery from TAM/vehicle injections, TAM-injected females weighed 1.16-fold more than vehicle injected females (17.88 ± .41 g veh vs. 20.68 ± 0.13 g TAM) (Fig 4B). TAM and vehicle-injected body weights did not significantly differ among male mice after 21d recovery (23.40 ± 0.81 g veh vs. 24.25 ± 0.45 g TAM) (Fig 4C).

**Figure 4.**
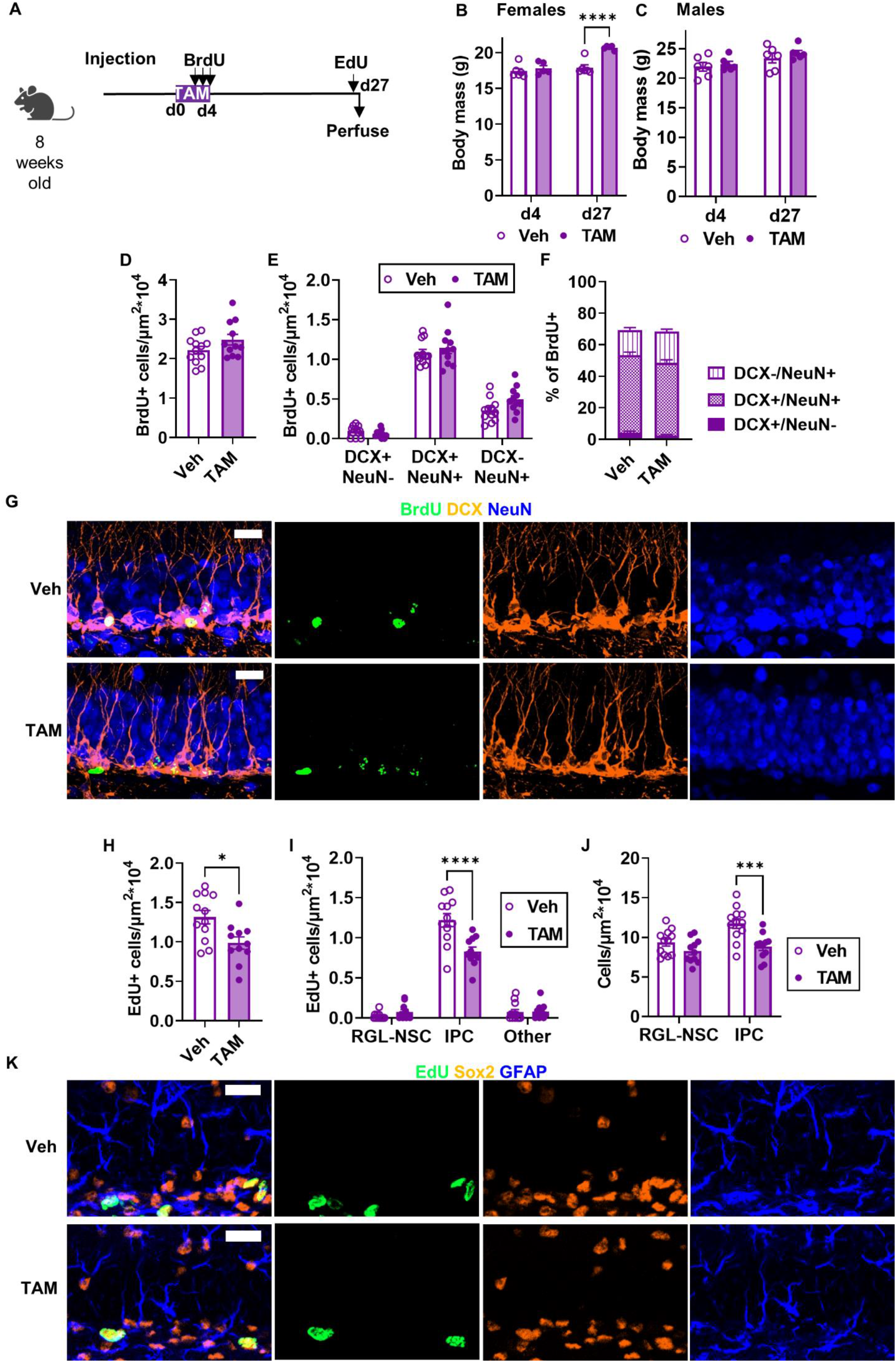
TAM injection suppresses progenitor cell proliferation 3 weeks after TAM. A) Experimental model and timeline. Adult wildtype mice were injected with TAM daily for 5 days then perfused 3 weeks later. Mice received 3 daily BrdU injections on the last 3 days of TAM and 1 EdU injection 2h before perfusion. B, C) Average body mass of females (2-way repeated measures ANOVA; day x TAM: *F*_*(1, 9)*_ = 11.17, *p* = 0.0086; day: *F*_*(1, 9)*_ = 22.53, *p* = 0.0010; TAM: F_*(1, 9)*_ = 21.55, *p* = 0.0012; subject: *F*_*(9, 9)*_ = 0.9448, *p* = 0.5330) and males (2-way repeated measures ANOVA; day x TAM: *F*_*(1, 10)*_ = 0.3575, *p* = 0.5632; day: F_(1, 10)_ = 24.83, *p* = 0.0006; TAM: *F*_*(1, 10)*_ = 0.6152, *p* = 0.4510; subject: *F*_*(10, 10)*_ = 6.138, *p* = 0.0042) at the end of TAM treatment (d4) and 3 weeks later (d27). Mean ± SEM of n = 5-6 female and 6 male mice. D) Density of BrdU+ cells (cells/μm^2^*10^4^) in the DG (unpaired t test, *t*_*(21)*_ = 1.613, *p* = 0.1217). Mean ± SEM of n = 11-12 mice. E) Density of DG BrdU+ cells co-labeled with DCX and/or NeuN (per μm^2^*10^4^) (2-way repeated measures ANOVA; cell type x TAM: *F*_*(2,42)*_ = 2.461, *p* = 0.0976; cell type: *F*_*(2, 42)*_ = 371.0, *p* < 0.0001; TAM: *F*_*(1, 21)*_ = 1.843, *p* = 0.1890; subject: *F*_*(21, 42)*_ = 1.698, *p* = 0.0714). Mean ± SEM of n = 11-12 mice. F) Percent of BrdU+ DG cells co-labeled for DCX and/or NeuN (2-way repeated measures ANOVA; cell type x TAM: *F*_*(2, 42)*_ = 3.027, *p* = 0.0591; TAM: *F*_*(1, 21)*_ = 0.1108, *p* = 0.7425; cell type: *F*_*(1*.*606, 33*.*73)*_ = 440.6, *p* < 0.0001; subject: *F*_*(21, 42)*_ = 0.5522, *p* = 0.9275). Mean ± SEM of n = 11-12 mice. G) Representative images of BrdU, DCX, and NeuN labeling. Scale = 20 μm. H) Density of EdU+ cells (cells/μm^2^*10^4^) in the DG (unpaired ttest, *t*_*(21)*_ = 2.814, *p* = 0.0104). Mean ± SEM of n = 11-12 mice. I) Density (cells/μm^2^*10^4^) of EdU+ phenotypic RGL-NSCs, IPCs, and other cells in the DG (2-way ANOVA; interaction: *F*_*(2, 63)*_ = 13.82, *p* < 0.0001; cell type: *F*_*(2, 63)*_ = 287.1, *p* < 0.0001; TAM: *F*_*(1, 63)*_ = 8.275, *p* = 0.0055). Mean ± SEM of n = 11-12 mice. J) Density (cells/ μm^2^*10^4^) of RGL-NSCs and IPCs in the DG (2-way repeated measures ANOVA; cell type x TAM: *F*_*(1, 21)*_ = 3.203, *p* = 0.0879; cell type: *F*_*(1, 21)*_ = 8.683, *p* = 0.0077; TAM: *F*_*(1, 21)*_ = 15.23, *p* = 0.0008; subject: *F*_*(21, 21)*_ = 1.029, *p* = 0.4741). Mean ± SEM of n = 11-12 mice. K) Representative images of EdU, SOX2, and GFAP labelling. Scale = 20 μm. \**P* < 0.05, \*\**P* < 0.01, \*\*\**P* < 0.001, \*\*\*\**P* < 0.0001, as determined by Sidak’s multiple comparisons test.

The total density of surviving BrdU+ cells did not differ between TAM and vehicle-injected mice (2.48 ± 0.14 *10^−4^ vs. 2.22 ± 0.10 *10^−4^ BrdU+ cells/µm^2^, respectively) (Fig 4D). The total density of BrdU+ cells expressing neuronal phenotypic markers DCX and/or NeuN also did not differ between TAM and vehicle-injected mice (Fig 4E,G). Almost half of BrdU+ cells in both groups co-expressed both DCX and NeuN (49.36 ± 1.84% veh and 46.24 ± 1.75% TAM) while slightly less than a fifth of BrdU+ cells in both groups were single positive for the mature neuronal marker NeuN (15.82 ± 1.47% veh and 19.78 ± 1.43% TAM) (Fig 4F,G). A small proportion of BrdU+ cells were single positive for DCX (4.12 ± 0.91% veh and 2.43 ± 0.55% TAM). Altogether, about 70% of BrdU cells showed a neuronal phenotype via DCX and/or NeuN co-labeling (69.38 ± 2.16% veh and 68.45 ± 1.71% TAM). Similar results were found when males and females were analyzed separately (Fig 5A-C). Combined, these results suggest that TAM injection during BrdU-labeling does not alter the number of new-born neurons 3 weeks later.

**Figure 5.**
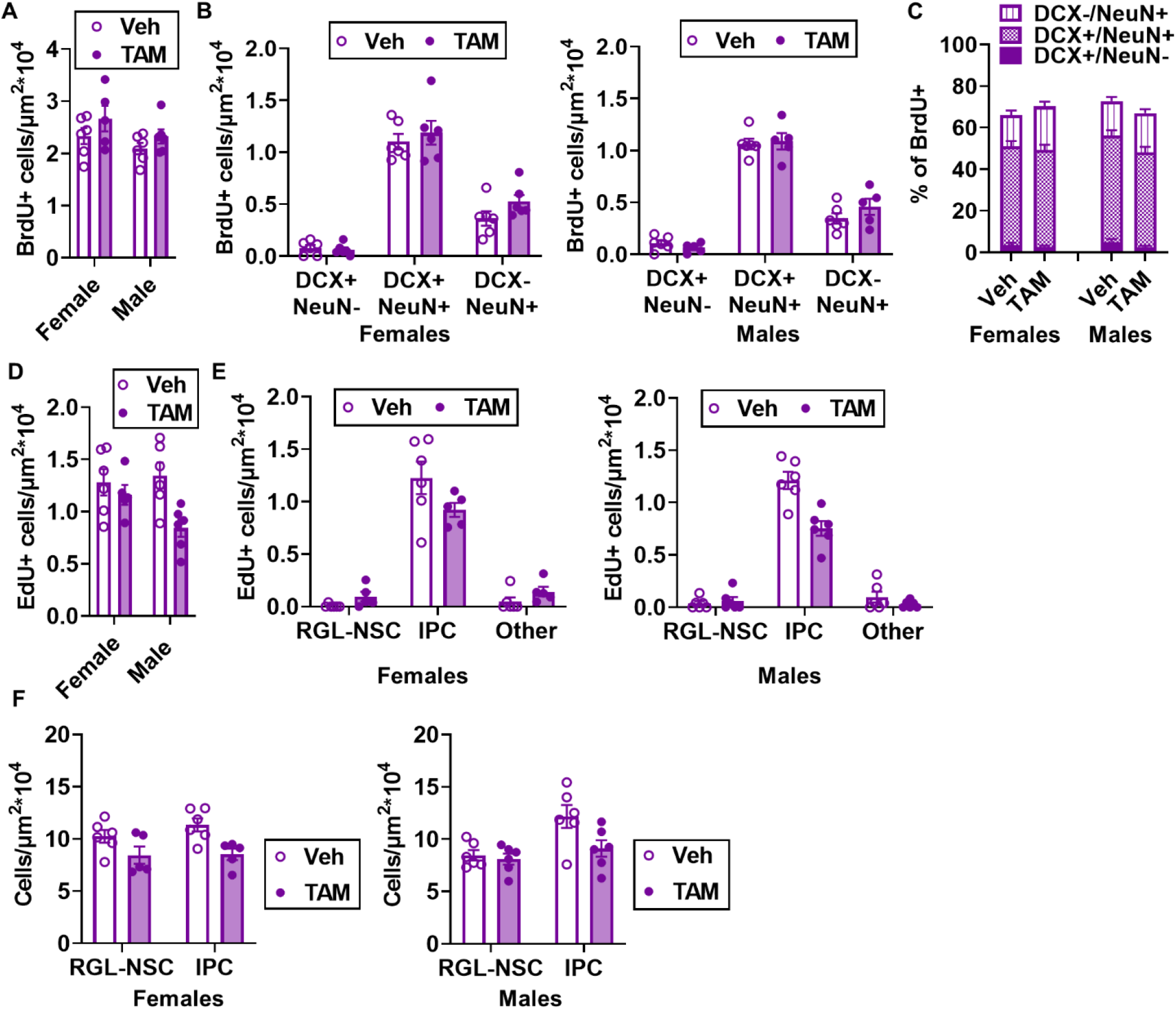
TAM injection effects on cell survival and proliferation do not differ by sex. A) Density of BrdU+ cells (cells/ μm^2^*10^4^) in the DG (2-way ANOVA; interaction: *F*_*(1, 19)*_ = 0.09813, *p* = 0.7575; sex: *F*_*(1, 19)*_ = 3.232, *p* = 0.0881; TAM: *F*_*(1, 19)*_ = 3.076, *p* = 0.0956). Mean ± SEM of n = 5-6 mice. B) Density of DG BrdU+ cells (cells/μm^2^*10^4^)co-labeled with DCX and/or NeuN (3-way repeated measures ANOVA; cell type: *F*_*(1*.*823, 34*.*63)*_ = 343.0, *p* < 0.0001; sex: *F*_*(1, 19)*_ = 0 5866, *p* = 0.4531; TAM: *F*_*(1, 19)*_ = 1.581, *p* = 0.2238; cell type x sex: *F*_*(2, 38)*_ = 0.5649, *p* = 0.5731; cell type x TAM: *F*_*(2, 38)*_ = 2.197, *p* = 0.1250; sex x TAM: *F*_*(1, 19)*_= 0.2991, *p* = 0.5908; cell type x sex x TAM: *F*_*(2, 38)*_ = 0.02294, *p* = 0.9773). Mean ± SEM of n = 6 female mice and 5-6 male mice. C) Percent of BrdU+ DG cells labeled for DCX and/or NeuN (3-way repeated measures ANOVA; cell type: *F*_*(1*.*608, 30*.*55)*_ = 401.7, *p* < 0.0001; sex: *F*_*(1, 19)*_ = 0.4138, *p* = 0.5277; TAM: *F*_*(1, 19)*_= 0.08557, *p* = 0.7731; cell type x sex: *F*_*(2, 38)*_ = 0.1847, *p* = 0.8321; cell type x TAM: *F*_*(2, 38)*_ = 2.828, *p* = 0.0716; sex x TAM: *F*_*(1, 19)*_ = 3.418, *p* = 0.0801; cell type x sex x TAM: *F*_*(2, 38)*_ = 0.04942), *p* = 0.9518. Mean ± SEM of n = 5-6 mice. D) Density of EdU+ cells (cells/μm^2^*10^4^) in the DG (2-way ANOVA; interaction: *F*_*(1, 19)*_ = 2.991, *p* = 0.0999; sex: *F*_*(1, 19)*_ = 1.313, *p* = 0.2660; TAM: *F*_*(1, 19)*_ = 7.928, *p* = 0.0110). Mean ± SEM of n = 5-6 mice. E) Density (cells/μm^2^*10^4^)of DG EdU+ phenotypic RGL-NSCs, IPCs, and other cells (3-way repeated measures ANOVA; cell type: *F*_*(2, 38)*_ = 274.4, *p* < 0.0001; sex: *F*_*(1, 19)*_ = 1.313, *p* = 0.2660; TAM: *F*_*(1, 19)*_ = 7.928, *p* = 0.0110; cell type x sex: *F*_*(2, 38)*_ = 0.4635, *p* = 0.6326; cell type x TAM: *F*_*(2, 38)*_ = 12.86, *p* < 0.0001; sex x TAM: *F*_*(1, 19)*_ = 2.991, *p* = 0.0999; cell type x sex x TAM: *F*_*(2, 38)*_ = 0.1659, *p* = 0.8477). Mean ± SEM of n = 5-6 female mice and 6 male mice. F) Density (cells/μm^2^*10^4^)of RGL-NSCs and IPCs in the DG, by sex (3-way repeated measures ANOVA; cell type: *F*_*(1, 19)*_ = 9.308, *p* = 0.0066; sex: *F*_*(1, 19)*_ = 0.1083, *p* = 0.7457; TAM: *F*_*(1, 19)*_ = 14.12, *p* = 0.0013; cell type x sex: *F*_*(1, 19)*_= 3.346, *p* = 0.0831; cell type x TAM: *F*_*(1, 19)*_ = 3.699, *p* = 0.0696; sex x TAM: *F*_*(1, 19)*_ = 0.3142, *p* = 0.5817; cell type x sex x TAM: *F*_*(1, 19)*_ = 0.7541, *p* = 0.3960). Mean ± SEM of n = 5-6 female mice and 6 male mice. \**P* < 0.05, \*\**P* < 0.01, \*\*\**P* < 0.001, \*\*\*\**P* < 0.0001, as determined by Sidak’s multiple comparisons test.

We also quantified cell proliferation of RGL-NSCs and IPCs 3 weeks after TAM using EdU to label proliferating cells. TAM injection led to a 1.32-fold suppression of total EdU+ cell density in the DG compared to vehicle-injected mice (1.31 ± 0.09 *10^−4^ cells/µm^2^ veh vs 0.99 ± 0.08 *10^−4^ cells/µm^2^ TAM) (Fig 4H, K). Classification of EdU+ cells as RGL-NSCs, IPCs or neither revealed that this reduction in EdU labeling was driven primarily by loss of EdU+ IPCs (Fig 4I, K). Quantification of total RGL-NSCs and IPCs similarly showed a significant loss of total IPC density in TAM-treated mice, with a more moderate, non-significant decrease in RGL-NSC density (Fig 4J, K). Similar results were found when males and females were analyzed separately (Fig 5D-F). These findings suggest that TAM injection causes a long-term suppression of IPC proliferation that is evident 3 weeks after TAM has ended.

### TAM chow enhances adult neurogenesis acutely but does not suppress cell proliferation long-term

To determine whether TAM chow alters adult neurogenesis, we fed adult wildtype C57BL6/J mice TAM or vehicle-matched chow for 14d, the last 3 of which were coupled with once per day BrdU injections to label dividing cells (Fig 6A). Mice were perfused 35d after the beginning of chow treatment (21d after the last TAM/BrdU injections), 2h after a single EdU injection to label acutely proliferating cells. Monitoring of food consumption confirmed that during the first ∼3d of chow exposure, mice consumed significantly less TAM chow than vehicle chow (2.88 ± 0.10g veh chow/ms/d vs 1.15 ± 0.15g TAM chow/ms/d), but consumption returned closer to vehicle chow levels shortly thereafter (Fig 6B; Fig 7A). Total average TAM consumption per mouse was 14.22 ± 0.95 mg, similar to that seen in Fig 1E (Fig 6B; Fig 7B). As expected from the TAM chow avoidance, mice that received TAM chow weighed less than vehicle chow fed mice at the end of chow exposure, though this pattern only reached significance in the male mice (Female: 16.65 ± 0.26g veh chow vs. 15.52 ± 0.31g TAM chow; Male: 22.37 ± 0.41g veh chow vs. 18.50 ± 0.45g TAM chow) (Fig 6C,D). In females, this pattern reversed after 21d recovery and by d35, TAM chow-fed mice weighed significantly more than vehicle chow-fed female mice (18.13 ± 0.25g veh chow vs. 20.00 ± 0.78g TAM chow) (Fig 6C). TAM and vehicle-fed male mice, in contrast, no longer differed in body weight after 3 weeks of recovery (24.07 ± 0.32g veh chow vs. 24.18 ± 0.88g TAM mice) (Fig 6D).

**Figure 6.**
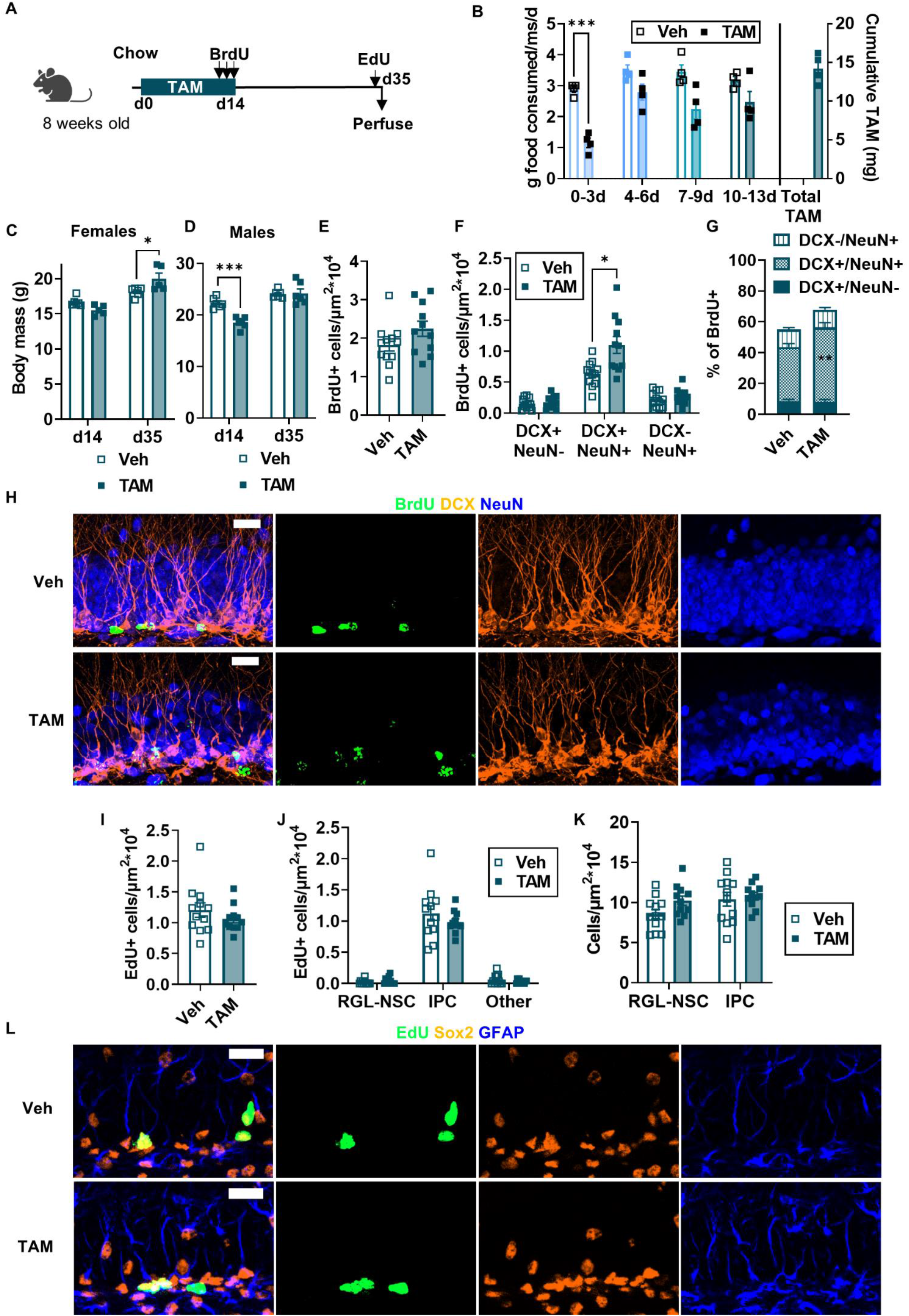
TAM chow alters the phenotype of surviving cells 35 days after initial exposure. A) Experimental model and timeline. Mice were injected with BrdU in the last 3 days of the 14-day TAM chow period. EdU was injected 35 days after initial chow exposure. B) Grams of food consumed per mouse per day over each 3-4 day period of diet exposure and TAM consumed in milligrams (2-way repeated measures ANOVA; day x TAM: *F*_*(3, 18)*_ = 2.322, *p* = 0.1095; day: *F*_*(1*.*564, 9*.*384)*_ = 8.978, *p* = 0.0088; TAM: *F*_*(1, 6)*_ = 45.86, *p* = 0.0005; subject: *F*_*(6, 18)*_ = 0.9896, *p* = 0.4613). Mean ± SEM of n = 4 mice. C) In female mice, body mass in grams 14d and 35d after diet initiation (2-way repeated measures ANOVA; day x TAM: *F*_*(1, 9)*_ = 10.62, *p* = 0.0099; day: *F*_*(1, 9)*_ = 42.05, *p* = 0.0001; TAM: *F*_*(1, 9)*_ = 0.8454, *p* = 0.3818; subject: *F*_*(9, 9)*_ = 0.7591, *p* = 0.6560). Mean ± SEM of n = 5-6 mice. D) In male mice, body mass in grams 14d and 35d after diet initiation (2-way repeated measures ANOVA; day x TAM: *F*_*(1, 10)*_ = 18.18, *p* = 0.0017; day: *F*_*(1, 10)*_ = 62.45, *p* < 0.0001; TAM: *F*_*(1, 10)*_ = 8.540, *p* = 0.0152; subject: *F*_*(10, 10)*_ = 1.886, *p* = 0.1658). Mean ± SEM of n = 6 mice. E) Density of BrdU+ cells (cells/μm^2^*10^4^) in the DG (unpaired t test, *t*_*(21)*_ = 1.675, *p* = 0.1087). Mean ± SEM of n = 11-12 mice. F) Density of DG BrdU+ cells (cells/μm^2^*10^4^) co-labeled with DCX and/or NeuN (2-way repeated measures ANOVA; cell type x TAM: *F*_*(2, 42)*_ = 8.654, *p* = 0.0007; cell type: *F*_*(1*.*207*_,_*25*.*35)*_ = 83.41, p < 0.0001; TAM: *F*_*(1, 21)*_ = 8.675, *p* = 0.0077; subject: *F*_*(21, 42)*_ = 1.516, *p* = 0.1239). Mean ± SEM of n = 11-12 mice. G) Percent of BrdU+ DG cells labeled for DCX and/or NeuN (2-way repeated measures ANOVA; TAM x cell type: *F*_*(2, 42)*_ = 6.501, *p* = 0.0035; TAM: *F*_*(1, 21)*_ = 16.13, *p* = 0.0006; cell type: *F*_*(1*.*545, 32*.*44)*_ = 151.5, *p* < 0.0001; subject: *F*_*(21, 42)*_ = 0.37 89, *p* = 0.9903). Mean ± SEM of n = 11-12 mice. H) Representative images of BrdU, DCX, and NeuN labeling. Scale = 20 μm. I) Density of EdU+ cells (cells/μm^2^*10^4^) in the DG (unpaired t test, *t*_*(21)*_= 1.043, *p* = 0.3090). Mean ± SEM of n = 11-12 mice. J) Density (cells/μm^2^*10^4^)of DG EdU+ phenotypic RGL-NSCs, IPCs, and other cells (2-way repeated measures ANOVA; cell type x TAM: *F*_*(2, 42)*_ = 1.157, *p* = 0.3241; cell type: *F*_*(1*.*066, 22*.*38)*_ = 210.3, *p* < 0.0001; TAM: *F*_*(1, 21)*_ = 1.087,*p* = 0.3090; subject: *F*_*(21, 42)*_ = 0.9645, *p* = 0.5209). Mean ± SEM of n = 11-12 mice. K) Density (cells/μm^2^*10^4^)of RGL-NSCs and IPCs in the DG (2-way repeated measures ANOVA; cell type x TAM: *F*_*(1, 21)*_ = 1.042, *p* = 0.3191; cell type: *F*_*(1, 21)*_ *-* 3.794, *p* = 0.0649; TAM: *F*_*(1, 21)*_ = 2.404, *p* = 0.1360; subject: *F*_*(21, 21)*_ = 1.138, *p* = 0.3846). Mean ± SEM of n = 11-12 mice. L) Representative images of EdU, SOX2, and GFAP labeling. Scale = 20 μm. \**P* < 0.05, \*\**P*< 0.01, \*\*\**p*< 0.001, \*\*\*\**P*< 0 0001, as determined by Sidak’s multiple comparisons test.

**Figure 7.**
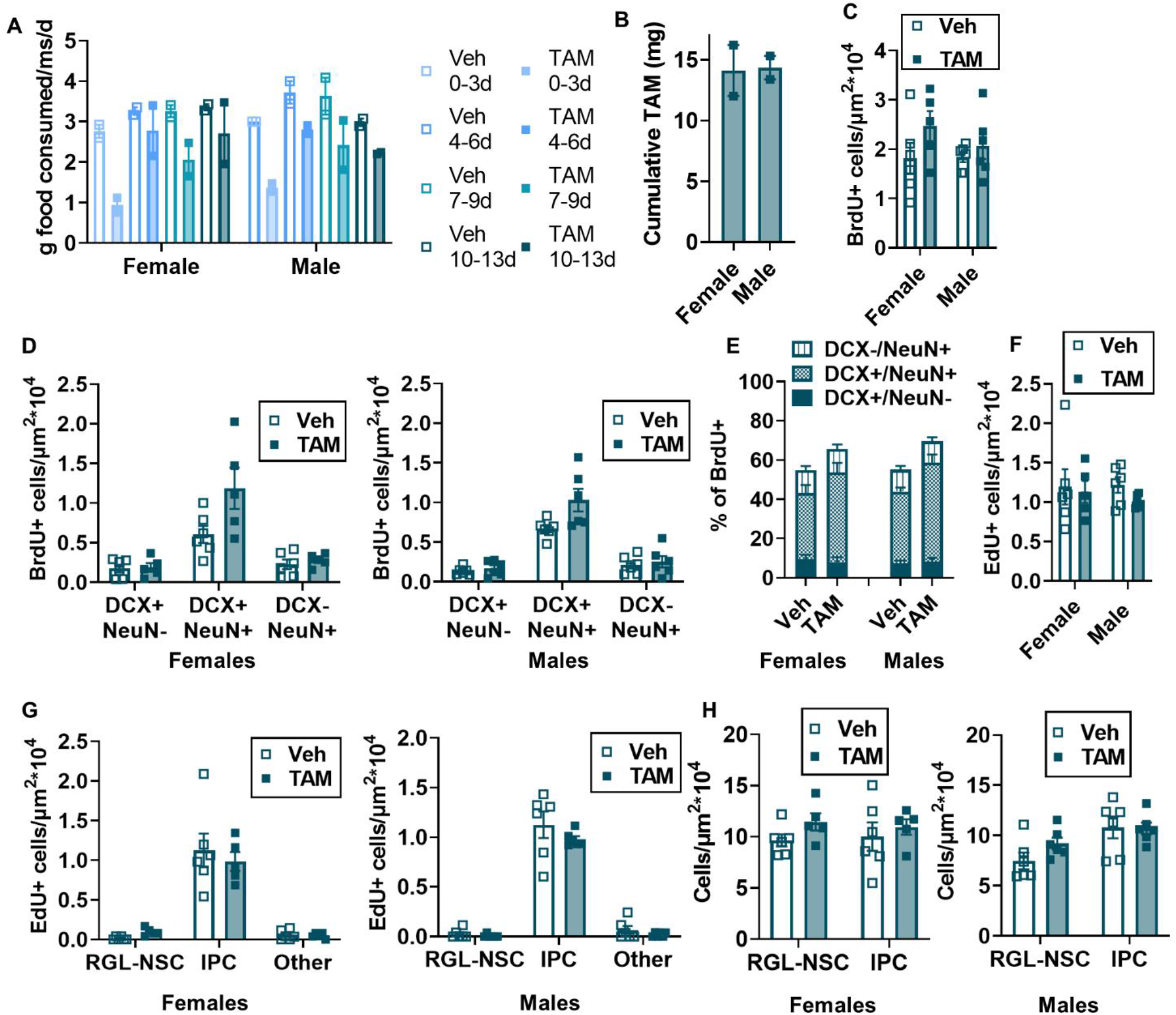
TAM diet effects on cell survival and proliferation do not differ by sex. A) Grams of TAM and vehicle chow consumed by male (n = 2 cages/sex) and female (n = 2 cages/sex) mice per mouse per day over each 3-4 day period of diet exposure. B) Cumulative TAM consumed in milligrams by sex (n = 2 cages/sex). C) Density of BrdU+ cells (cells/μm^2^*10^4^) in the DG, by sex (2-way ANOVA; interaction: *F*_*(1, 19)*_ = 0.6818, *p* = 0.4192; sex: *F*_*(1, 19)*_ = 0.5413, *p* = 0.4709; TAM: *F*_*(1, 19)*_ = 2.921, *p* = 0.1037). Mean ± SEM of n = 5-6 mice. D) Density of DG BrdU+ cells (cells/μm^2^*10^4^)co-labeled with DCX and/or NeuN in females and males (3-way repeated measures ANOVA : cell type: *F*_*(1*.*213, 23*.*05)*_= 78.09, *p* < 0.0001; sex: *F*_*(1, 19)*_ = 0.2586, *p* = 0.6169; TAM: *F*_*(1, 19)*_ = 8.323, *p* = 0.0095; cell type x sex: *F*_*(2, 38)*_ = 0.02142, *p* = 0.9788; cell type x TAM: *F*_*(2, 38)*_ = 8.256, *p* = 0.0011; sex x TAM: *F*_*(1, 19)*_ = 0.3225, *p* = 0.5767; cell type x sex x TAM: *F*_*(2, 38)*_ = 0.5548, *p* = 0.5787). Mean ± SEM of n = 5-6 female mice and 6 male mice. E) Percent of BrdU+ DG cells labeled for DCX and/or NeuN in male and female mice (3-way repeated measures ANOVA : cell type: *F*_*(1*.*557, 29*.*58)*_ = 140.5, *p* < 0.0001; sex: *F*_*(1, 19)*_ = 0.3592, *p* = 0.5561; TAM: *F*_*(1, 19)*_ = 14.62, *p* = 0.0011; cell type x sex: *F*_*(2, 38)*_ = 0.6323, *p* = 0.5369; cell type x TAM: *F*_*(2, 38)*_ = 5.877, *p* = 0.0060; sex x TAM: *F*_*(1, 19)*_ = 0.3233, *p* = 0.5763; cell type x sex x TAM: *F*_*(2, 38)*_ = 0.06331, *p* = 0.9387). Mean ± SEM of n = 5-6 mice. F) Density of EdU+ cells (cells/ μm^2^*10^4^)in the DG in male and female mice (2-way ANOVA; interaction: *F*_*(1, 19)*_ = 0.2581, *p* = 0.6173; sex: *F*_*(1, 19)*_ = 0.1190, *p* = 0.7339; TAM: *F*_*(1, 19)*_ = 0.9220, *p* = 0.3490). Mean ± SEM of n = 5-6 mice. G) Density (cells/ μm^2^*10^4^) of DG EdU+ phenotypic RGL-NSCs, IPCs, and other cells in male and female mice (3-way ANOVA : cell type: *F*_*(1*.*058, 20*.*10)*_ = 190.3, *p* < 0.0001; sex: *F*_*(1, 19)*_ = 0.1190, *p* = 0.7339; TAM: *F*_*(1, 19)*_ = 0.9220, *p* = 0.3490; cell type x sex: *F*_*(2, 38)*_ = 0.05511, *p* = 0.9465; cell type x TAM: *F*_*(2, 38)*_ = 1.094, *p* = 0.3451; sex x TAM: *F*_*(1, 19)*_ = 0.2581, *p* = 0.6173; cell type x sex x TAM: *F*_*(2, 38)*_ = 0.08267, *p* = 0.9208). Mean ± SEM of n = 5-6 female mice and 6 male mice. H) Density (cells/ μm^2^*10^4^) of RGL-NSCs and IPCs in the DG of male and female mice (3-way ANOVA: cell type: *F*_*(1, 19)*_ = 3.943, *p* = 0.0617; sex: *F*_*(1, 19)*_ = 2.181, *p* = 0.1561; TAM: *F*_*(1, 19)*_ = 2.703, *p* = 0.1166; cell type x sex: *F*_*(1, 19)*_ = 4.370, *p* = 0.0503; cell type x TAM: *F*_*(1, 19)*_ = 1.338, *p* = 0.2618; sex x TAM: *F*_*(1, 19)*_ = 0.1703, *p* = 0.6845; cell type x sex x TAM: *F*_*(1, 19)*_ = 0.1708, *p* = 0.6840). Mean ± SEM of n = 5-6 female mice and 6 male mice. \**P*< 0.05, \*\**P*< 0.01, \*\*\**P*< 0.001, \*\*\*\**P*<0.0001, as determined by Sidak’s multiple comparisons test.

The total density of surviving BrdU+ cells did not significantly differ between TAM and vehicle chow-fed mice (2.25 ± 0.20 *10^−4^ cells/µm^2^ vs 1.83 ± 0.16 *10^−4^ cells/µm^2^, respectively) (Fig 6E). In contrast, the density of BrdU+ cells co-expressing both DCX and NeuN was significantly higher in TAM chow-fed mice than vehicle-fed mice (0.64 ± 0.06 *10^−4^ cells/µm^2^ veh chow vs. 1.10 ± 0.14 *10^−4^ cells/µm^2^ TAM chow) (Fig 6F,H). The percent of BrdU+ cells co-expressing both DCX and NeuN was similarly higher in the TAM chow-fed mice than vehicle chow-fed mice (35.15 ± 2.17% veh chow vs. 48.09 ± 3.42% TAM chow) (Fig 6G). The percentage of BrdU+ cells single positive for DCX or NeuN was small in both groups and did not differ between groups (DCX: 8.46 ± 1.26% veh chow vs. 7.915 ± 1.63% TAM chow; NeuN: 11.38 ± 1.31% veh chow vs. 11.42 ± 1.57% TAM chow). Similar results were found when males and females were analyzed separately (Fig 7C-E). These results suggest that the TAM chow exposure paradigm drives more neurogenesis primarily via enhanced neuronal differentiation of cells proliferating during TAM exposure.

We also quantified cell proliferation of RGL-NSCs and IPCs 3 weeks after TAM using EdU to label proliferating cells. EdU+ cell density in the DG did not significantly differ between veh and TAM chow fed mice (1.21 ± 0.12 *10^−4^ cells/µm^2^ veh vs 1.06 ± 0.07 *10^−4^ cells/µm^2^ TAM) (Fig 6I, L). Classification of EdU+ cells as RGL-NSCs, IPCs or neither similarly showed no difference in EdU+ density of these cell subtypes between chow groups (Fig 6J, L). Quantification of total RGL-NSCs and IPCs also showed no difference in total RGL-NSC or IPC density (Fig 6K, L). Similar results were found when males and females were analyzed separately (Fig 7F-H). These findings suggest that TAM chow feeding does not strongly alter cell proliferation relative to vehicle chow 3 weeks after chow treatment has ended.

## Discussion

In this work, we found that TAM chow treated mice showed less recombination efficiency but similar recombination specificity as TAM injected mice in a commonly used NestinCreER^T2^ model that targets recombination in adult NSPCs. Both administration protocols also had inherent, though different, effects on adult neurogenesis. The TAM injection protocol suppressed IPC proliferation 3 weeks after TAM, whereas the TAM chow protocol did not disrupt later IPC proliferation but did induce greater neuronal differentiation of cells born during TAM. Altogether, our results suggest that both administration routes of TAM are potentially useful but selection of controls requires attention to the inherent effects of TAM delivery.

The difference between TAM injection and TAM chow in recombination efficiency is most likely explained by total TAM exposure. Recombination efficiency in tamoxifen-inducible transgenic systems is partly dose-dependent (Hayashi and McMahon, 2002) and mice given TAM chow were exposed to overall less TAM than those given TAM injections. Mice showed strong aversion to the TAM chow, as evidenced by the ∼1 week delay between chow introduction and substantial daily consumption. The chow was sweetened to increase palatability and delivered on the cage floor in dishes to ease access, so future efforts to overcome this barrier to TAM ingestion may require longer TAM chow treatments. In studies targeting adult neurogenic processes, however, longer chow treatment will result in a more heterogenous population of recombined cells at different stages of differentiation. Temporal precision is often a goal of adult neurogenesis studies, and extending chow treatment times may not always fit with that goal. Another possibility would be to provide TAM via oral gavage. However, oral gavage is notoriously stressful for animals (Brown et al., 2000) and requires expertise on the part of the experimenter to reduce that stress and avoid injury to animals (Arantes-Rodrigues et al., 2012). If high recombination efficiency and selective targeting of cells of similar maturity are desired, our data suggest that TAM injections are still the preferable method.

Despite differences in recombination efficiency between the TAM administration routes tested here, this work also shows that 2 weeks of TAM chow induces genetic recombination with similar specificity to 5d of TAM injection. Mice fed TAM chow for 2 weeks showed a recombined cell population of similar phenotype (primarily NSPCs) as TAM injected mice 7d after the start of injections. Thus, TAM-infused chow could be a workable alternative administration route when TAM injection is not feasible, when lower recombination efficiency is desired, or when long-term suppression of cell proliferation would interfere with interpretation of experimental results.

Our findings comparing TAM versus vehicle injected mice provide several useful insights into the effects of this commonly-used agent on adult neurogenesis. First, we found that neurogenesis derived from cells born during TAM was not substantially affected by TAM injection. These results are similar to those reported in (Rotheneichner et al., 2017), which showed no difference in BrdU+ cell number or phenotype between TAM and vehicle injected mice 10 days after TAM/BrdU using 5 month old male and female mice. In contrast, using 3-4 week old (juvenile) male mice, Lee et al. (2020) found that the number of surviving BrdU+ cells ∼1 week after TAM/BrdU was suppressed by ∼2 fold in TAM injected versus vehicle injected mice. Combined, these findings suggest that the effect of TAM injection on production of new DG cells that survive 1+ weeks could depend on organismal age. Sex may also be a factor, but this is difficult to discern as previous work either used only males or did not show data for males and females separately. Future research is needed to address the effects of TAM on cell proliferation and survival to better understand how TAM affects neurogenesis across the lifespan and in both sexes.

Second, we show that TAM injection had prolonged effects on progenitor cell proliferation, suppressing progenitor number and proliferation by ∼1.3 fold 3 weeks after TAM. Lee et al. similarly reported a ∼1.8 fold TAM-induced suppression of Mki67-labeled progenitors one week after TAM in juvenile mice. Rotheneicher et al. reported no difference in PCNA+ progenitor cell number 10 days after TAM in their 5 month old mice, however. Again, this difference may reflect increased susceptibility of juvenile mice to the effects of TAM, a subtle sex difference, or a timepoint effect.

The mechanism by which TAM suppresses progenitor proliferation weeks after TAM treatment is unclear. One possible mechanism for suppression of cell proliferation associated with TAM treatment in general is brain estrogen receptor modulation by TAM. TAM acts on brain ERα, ERβ, and the transmembrane receptor GPR30 (Gonzalez et al., 2016). Specifically, TAM can be an estrogen agonist at GPR30 (Filardo et al., 2000; Vivacqua et al., 2006a, 2006b) and Erβ (McDonnell and Wardell, 2010), and it can induce estrogen blockade at ERα (McDonnell and Wardell, 2010). Because ERα agonists can enhance cell proliferation in the adult hippocampus (Mazzucco et al., 2006), it therefore seems possible that TAM blockade of ERα could suppress proliferation. Receptor-independent effects on cell cycle genes, as proposed by Lee et al. 2020, are also a potential mechanism of TAM-induced suppression of proliferation. With either mechanism, though, it is unclear why those effects would persist long after TAM withdrawal. TAM metabolites remain in the brain for about 8 days after administration (Valny et al., 2016), making it unlikely that suppression of proliferation 3 weeks later is a result of active TAM presence. This long-term persistence could therefore be because initial effects of TAM are slow to arise and/or indelible (e.g. epigenetic changes), or because they reflect effects of TAM withdrawal rather than TAM itself. Future research is needed to parse out these multiple, non-exclusive hypotheses.

We also noted delayed weight gain in female mice that received TAM injection, resulting in higher average body mass of TAM treated females than vehicle treated females 3 weeks after injection. Because TAM was administered systemically, this delayed weight gain in female mice could be due to TAM effects on multiple central or peripheral tissues. One candidate tissue is the hypothalamus. In mice, hypothalamic ERα-dependent gene expression changes have been linked to common TAM side effects, including decreased movement (Zhang et al., 2021), which could contribute to weight gain.

TAM chow had qualitatively different effects on neurogenesis and cell proliferation than TAM injection. In contrast to TAM injections, TAM chow increased neuronal differentiation of cells born during TAM but had no effect on density of EdU+ cells 3 weeks after TAM. Caloric restriction and reduced total TAM exposure in chow fed mice (compared with TAM injected mice) both likely contribute to this difference between paradigms. Caloric restriction is a particularly important confounding factor to consider when comparing TAM chow-fed to vehicle chow-fed animals. Though Bondolfi et al. found no effect of caloric restriction on hippocampal neurogenesis in male mice aged 3-11 months (Bondolfi et al., 2004), others have found that dietary restriction and intermittent fasting enhance neuronal differentiation in the adult hippocampus (Hornsby et al., 2016; Li et al., 2020). Both the caloric restriction during TAM feeding and the return to normal feeding after the end of TAM chow may therefore be affecting neurogenesis in TAM-fed mice. Regardless of the reason for differing effects of TAM by administration route, our data further underscore the need for internal TAM-treated controls in experiments using TAM chow, just as with TAM injection.

### Limitations and Future Directions

Because this work examines direct TAM effects on experimental endpoints only in the context of neurogenesis research in adult animals, further testing is needed to determine the suitability of alternate TAM dosing methods for subadult animals and research on other brain processes. Also, our BrdU+ counts reflect both cell proliferation during TAM administration and post-TAM cell survival, making it difficult to disentangle TAM effects on either process independently. Better understanding of the molecular mechanism by which TAM acts on adult neurogenesis, when provided by injection or chow, is also still needed. Finally, clarity regarding the actions of TAM on brain estrogen receptors is lacking and could help guide researchers’ choice of TAM dosing methods and interpretation of studies where TAM is used.

### Conclusions and Recommendations

This work shows that voluntary TAM chow consumption may be a suitable alternative to TAM injections for inducing Cre-lox recombination in some adult neurogenesis studies. This work further identifies several effects of TAM administration protocols, whether by injection or food, on adult neurogenesis endpoints. These effects are separate from genetic recombination effects and are an important confounding variable in experimental designs that rely on comparisons to a vehicle-treated control. Thus, we suggest that research using TAM-inducible Cre lines use TAM-treated wildtype (non-recombination susceptible) littermates as controls, rather than vehicle-treated mice.

## Materials and methods

### Mice

All mice were group housed with 3-5 mice per cage in The Ohio State University Psychology building mouse vivarium in standard ventilated cages on a 12/12 h light/dark cycle (lights on 6:30 A.M.) with *ad libitum* access to food and water. Male and female mice were eight to nine weeks old at the time of the experiment and housed in groups of two to four. All animal use was in accordance with institutional guidelines approved by The Ohio State University Institutional Animal Care and Use Committee.

### Experimental Design and Statistical Analysis

#### Study 1: TAM injection versus TAM chow

7-9 week old NestinCreER^T2^ (Lagace et al., 2007)(Jackson #016261);R26R-LoxP-STOP-LoxP-EYFP (Srinivas et al., 2001) (Jackson #007909) littermates were randomly assigned by whole cage to one of 4 groups: TAM inject d7 perfuse (n = 10), TAM inject d14 perfuse (n = 11), TAM chow 10d (n = 7), TAM chow 14d (n = 11). TAM injected mice received daily 180 mg/kg/day IP injections of TAM for 5 days. TAM chow mice were first acclimated to standard chow provision in a dish on the cage floor for 3 days before switching to TAM chow. Food was weighed and refreshed every 2-4 days. At the indicated tissue harvest time, mice were anesthetized with an 87.5 mg/kg ketamine, 12.5 mg/kg xylazine mixture and then transcardially perfused with ice-cold 0.1 M PBS. Body weight data were analyzed by a repeated measures mixed-effects model (Day x TAM) within each sex, followed by Sidak’s multiple comparisons within day. Food consumption was analyzed by two-way ANOVA (sex x day) followed by Sidak’s multiple comparisons between days within sex. Cell count data were analyzed by one-way ANOVA followed by Tukey’s posthoc tests or by two-way ANOVA (hippocampal subregion x group) followed by Tukey’s posthoc multiple comparisons between groups within subregion.

#### Study 2: TAM versus vehicle injection

Wildtype C57Bl6/J mice were obtained at 6 weeks of age from Jackson Labs (#000664) and allowed to acclimate for 2 weeks. Mice were assigned randomly by whole cage to either TAM (n = 11) or vehicle (sunflower oil, n = 12) injection groups. TAM injections were as described for study 1. On the last 3 days of TAM/vehicle injection, mice also all received daily BrdU injections (IP, 150 mg/kg/d). 21d after the last TAM/BrdU injections, mice received a single injection of EdU (IP, 150 mg/kg) and were perfused as in Study 1 2h later. Body weight data were analyzed by a repeated measures mixed-effects model (Day x TAM) within each sex, followed by Sidak’s multiple comparisons within day. BrdU+/EdU+ density was compared by unpaired t-test. Density and proportion of BrdU/DCX/NeuN or EdU/GFAP/SOX2 cell subtypes was compared by two-way repeated measures ANOVA followed by posthoc Sidak’s multiple comparisons within cell type.

#### Study 3: TAM versus vehicle chow

Wildtype C57Bl6/J mice were obtained at 6 weeks of age from Jackson Labs (#000664) and allowed to acclimate for 1 week. Mice were assigned randomly by whole cage to either TAM (n = 11) or vehicle (n = 12) chow groups. Diet was provided as described for Study 1 for 14d. On the last 3 days of TAM/vehicle chow, mice also all received daily BrdU injections (IP, 150 mg/kg/d). 21d after the last TAM/BrdU injections, mice received a single injection of EdU (IP, 150 mg/kg) and were perfused as in Study 1 2h later. Food consumption was analyzed by repeated measures two-way ANOVA (chow type x day) followed by Sidak’s multiple comparisons between chow type within days. Remaining analysis for this experiment is similar to that for study 2.

### TAM injection preparation

TAM (Fisher, #50-115-2413) was dissolved overnight at 20 mg/ml at 37°C in sterile sunflower oil then stored at +4°C for 1 week maximum, all while protected from light.

### TAM and vehicle chow

TAM chow (500 mg TAM/kg diet, TD.130858, contains 49.5 g/kg sucrose) and matched vehicle chow were obtained from Envigo and stored at +4°C until given to mice.

### BrdU and EdU preparation

BrdU (Sigma, #B5002) and EdU (Click Chemistry Tools, #1149-500) were each dissolved fresh in sterile physiological saline at 10 mg/ml.

### Tissue processing

Tissue fixation, slicing, and immunofluorescent staining were performed similarly to our previous work (Dause and Kirby, 2020) using the antibodies described in Table 1. In brief, brains were post-fixed in 4% paraformaldehyde (Fisher #AC169650010) in 0.1M phosphate buffer at +4°C for 24h then equilibrated in 30% sucrose (Fisher S5-3) in 0.1M phosphate buffered saline (PBS, Fisher #BP399-20) at +4°C before slicing on a freezing microtome (Leica) in a 1:12 series of 40 µm thick coronal sections. Sections were stored in cryoprotectant medium at -20°C until processed for immunofluorescent staining. Free-floating sections were rinsed in PBS, blocked in 1% normal donkey serum (Jackson Immunoresearch #017000121), 0.3% triton-100X (Fisher #AC215682500) in PBS and incubated in primary antibody in blocking solution overnight at 4°C on rotation. The next day, sections were rinsed, incubated in secondary antibody in blocking solution for 2h at room temperature on rotation and then rinsed and counterstained with Hoechst 33342 (1:2000 in PBS, Fisher #H3570) before being mounted on SuperFrost Plus slides (Fisher #12-550-15), dried and coverslipped with Prolong Gold Antifade mounting medium (Fisher #P36934). For BrdU staining, sections were processed for other antibodies first then postfixed fixed in 4% paraformaldehyde for 10 min at room temperature. Sections were then rinsed and incubated in 2N HCl (Fisher A144500) at 37°C for 30 min, followed by rinsing, blocking and primary/secondary incubation as described above. For EdU click labeling, sections were click reacted using a Click&Go EdU 488 imaging kit (Click Chemistry Tools, #1324) according to manufacturer instructions before proceeding with subsequent immunolabeling. Slides were all dried overnight at room temperature in the dark and then stored long-term at +4 °C.

**Table 1:** Antibodies used in immunohistochemical staining.

| Table 1: Antibodies used in immunohistochemical staining. |  |  |  |  |  |
| --- | --- | --- | --- | --- | --- |
| Primary antibody | Vendor/product no. | Dilution | Secondary | Vendor/product no. | Dilution |
| Rabbit anti-GFP | ThermoFisher A11122 | 1:1000 | Donkey Alexafluor488 anti-rabbit | Fisher A21206 | 1:500 |
| Mouse anti-GFAP | EMD Millipore MAB360 | 1:1000 | Donkey Alexafluor647 anti-mouse | Fisher A31571 | 1:500 |
| Rat anti-SOX2 | Affymetrix eBioscience 14-9811 | 1:1000 | Donkey Alexafluor594 anti-rat | Fisher A21209 | 1:500 |
| Rabbit anti-DCX | Cell Signaling 4604 | 1:500 | Donkey Alexafluor555 anti-rabbit | Fisher A31572 | 1:500 |
| Mouse anti-NeuN | EMD Millipore MAB377 | 1:1000 | Donkey Alexafluor647 anti-mouse | Fisher A31571 | 1:500 |
| Rat anti-BrdU | BioRad OBT0030 | 1:500 | Donkey Alexafluor488 anti-mouse | Fisher A21202 | 1:500 |

### Cell quantification

Similar to our previous work (Dause and Kirby, 2020), the DG of the hippocampus was imaged in 15-μm z-stacks at 20× magnification using a Zeiss Axio Observer Z.1 with apotome digital imaging system and Axiocam 506 monochrome camera (Zeiss). EYFP+ cells were identified based on EYFP+ cytoplasm around a Hoechst+ nucleus. EYFP+ cells were considered RGL-NSCs if they had a SOX2+ nucleus in the SGZ and colocalized with GFAP in the cytoplasm with an apical morphology. They were considered IPCs if they had a SOX2+ nucleus in the SGZ but did not have a GFAP+ cytoplasm. They were considered astrocytes if they had a SOX2+ nucleus and colocalized with GFAP in the cytoplasm with a stellate morphology. They were considered immature neurons/neuroblasts if they had a Hoechst+ nucleus in the SGZ and a cytoplasm that co-localized with DCX. BrdU+ cells were counted in the SGZ and granule cell layer throughout the DG. Colabeling with DCX and NeuN was assessed in each BrdU+ cell in the z-stacks as surrounding DCX+ cytoplasm and/or nuclear NeuN overlap with BrdU. All cell counts were divided by DG area sampled to yield a density of cells.

